# Crystal structure reveals the full Ras:Raf interface and advances mechanistic understanding of Raf activation

**DOI:** 10.1101/2020.07.28.225938

**Authors:** Trinity Cookis, Carla Mattos

## Abstract

The interaction between Ras and Raf-kinase through the Ras-binding (RBD) and cysteine-rich domains (CRD) of Raf is essential for signaling through the mitogen-activated protein kinase (MAPK) pathway, yet the molecular mechanism leading to Raf activation has remained elusive. We present the 2.8 Å crystal structure of the HRas/CRaf-RBD_CRD complex showing the Ras/Raf interface as a continuous surface on Ras. In the Ras dimer, with helices roughly perpendicular to the membrane, the CRD is located between the two Ras protomers and far from the membrane, where its dynamic nature in the Ras binding pocket is expected to accommodate BRaf and CRaf heterodimers. Our structure and its analysis by MD simulations, combined with work in the literature, result in a molecular model in which Ras binding is involved in the release of Raf autoinhibition while the Ras/Raf complex dimerizes to promote a platform for signal amplification, with Raf-CRD poised to have direct and allosteric effects on both the Ras active site and the dimerization interface.

## Introduction

Ras interacts with Raf kinase through Raf’s N-terminal Ras-binding (RBD) and cysteine-rich (CRD) domains to drive cell proliferation through the Ras/Raf/MEK/ERK mitogen activated protein kinase (MAPK) pathway (*1-3*). Mutations in Ras and Raf are major drivers of human cancers (*4, 5*) and in spite of great efforts over the last 20 years, the mechanism for Ras-mediated activation of Raf remains elusive. This critical gap in our basic understanding of the pathway has limited our ability to explore novel strategies to develop drugs against Ras-driven cancers (*6*).

We previously solved crystal structures for wildtype Ras and its Q61L mutant bound to the Raf-RBD, revealing that this low nanomolar affinity complex occurs through electrostatic interactions at switch I (residues 30-40 in Ras) and that the oncogenic mutant RasQ61L has global impacts on the dynamics of the Ras/Raf complex (*7-10*). The transformation potential of Raf, however, requires interaction with Ras through both the Raf-RBD and Raf-CRD (*2, 11, 12*), yet understanding of the weak micromolar affinity interaction between Ras and the Raf-CRD has remained limited due to lack of structural information. The Raf-CRD requires coordination of two zinc ions and has been shown to interact with various phospholipids and supported lipid bilayers *in vitro* (*3, 13-16*). Its membrane-binding role has further been studied by molecular dynamics (MD) simulations and recently supported by NMR guided characterization of nanodisc-bound KRas/CRaf-RBD_CRD complexes (*17-22*).

In addition to the N-terminal Ras-binding conserved region (CR1) (residues 52-194, CRaf numbering), the three Raf isoforms (ARaf, BRaf, and CRaf) share two additional regions with high similarity that are connected through unstructured linker regions (*23*). The second conserved region (CR2) (residues 254-269) contains a serine/threonine rich domain and the third conserved region (CR3) comprises the C-terminal kinase domain (residues 349-609) (Fig. 1A) (*24-26*). Prior to its recruitment to the membrane and interaction with Ras, Raf is held in an autoinhibited state through the aid of 14-3-3 scaffold proteins that bind motifs located in the CR2 and CR3 regions (*27-29*). Recruitment of Raf to the membrane through Ras results in dimerization of the Raf kinase domain, leading to its activation and subsequent phosphorylation of MEK to propagate signaling (*30, 31*). Recent cryo-EM structures of inactive and active BRaf/14-3-3 and BRaf/MEK1/14-3-3 complexes have shed light into this mechanism revealing a critical role for the Raf-CRD in stabilizing the autoinhibited state (*27, 32, 33*). However, Ras’ involvement in the release of Raf autoinhibition has not yet been mechanistically understood.

**Fig. 1.**
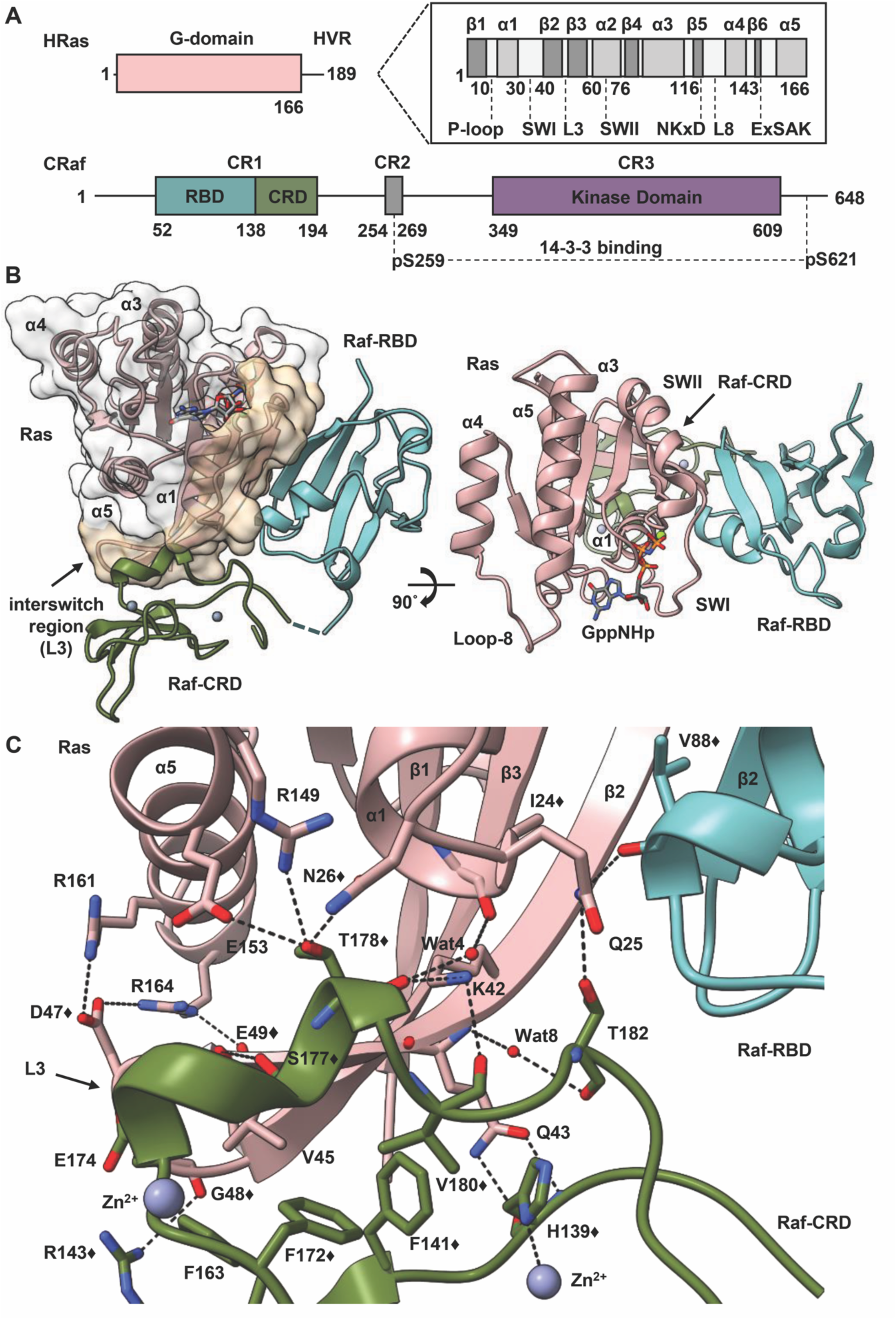
Structure of the HRas-CRaf-RBD_CRD complex. **(A)** Schematic diagrams of HRas and CRaf provide structural boundaries for the Ras G-domain and hypervariable region, and the three conserved regions found in Raf kinases. **(B)** Cartoon and surface representations of the HRas/CRaf-RBD_CRD complex reveal a continuous binding surface spanning Ras (pink) residues 23 through 48 for both the Raf-RBD (teal) and Raf-CRD (green). **(C)** The Raf-CRD interacts with residues in Ras helix 1 and the interswitch region through hydrogen bonds and hydrophobic contacts. The symbol ♦ identifies residues that show chemical shift perturbations in the analysis described by Feng et al. (*17*).

Dimerization of the Ras G-domain through an interface involving helices 4 and 5 is critical for signaling through the MAPK cascade (*34-37*) and we recently reported that Ras dimerization is promoted through the binding of the Raf-RBD (*38*). Here we present the crystal structure of HRas bound to a construct containing both the CRaf-RBD and CRD at 2.8 Å resolution (Fig. 1B-C & Table S1). This structure reveals that the Raf-CRD interacts with Ras residues at a surface contiguous to the Raf-RBD along beta 2, stretching to loop 3 at the interswitch region. From this structure we built a model of the Ras/Raf dimer, in which the Raf-CRD nestles at the base of the two Ras protomers, and is poised to have a significant impact on the Ras active site and beyond. Based on our crystal structure and recent developments in the literature, we propose a structural model for the role of Ras in the activation of Raf.

## Results

### Overall structure of Ras/Raf-RBD_CRD complex

Together, the Raf RBD and CRD binding interfaces are contiguous surfaces spanning Ras residues 23 through 48, including the C-terminal end of the Ras helix 1, switch I, beta 2 and loop 3 (Fig. 1B). The Raf-RBD interacts with Ras at switch I and the N-terminal portion of beta 2 (residues 27-41) as previously described (*7*) and the short linker connecting the Raf-RBD and Raf-CRD (residues 133-137) is partially disordered in our structure. The Raf-CRD interaction involves hydrogen bonds and hydrophobic contacts with C-terminal residues in Ras helix 1 (residues L23-N26) and beta strand 2, extending to loop 3 in the interswitch region (residues K42-G48), with direct and allosteric connections to helix 5 (Fig. 1C). The Raf-CRD interface involves residues near the CRD C-terminus in the stretch from F172 to T182, and nearby N-terminal residues H139, F141, R143, as well as F163. This interface includes the helix adjacent to the first Zn^2+^ binding site and extends toward the C-terminus, where the second Zn^2+^ ion is coordinated by residues C184 as well as by the interface residue H139, linking the N and C-terminal ends of the Raf-CRD. The position of the CRD observed in our crystal structure is consistent with recently published NMR chemical shift perturbation (CSP) data (Fig. 1C & S1A), and with Ras N26G and V45E mutations shown to disrupt the Ras/Raf-CRD interaction (*12, 17*). Given its close contact with loop 3 on Ras, the Raf-CRD is allosterically connected to helix 5 through salt bridges that form between Ras D47 and E49 on loop 3 and R161 and R164 on helix 5 (Fig. 1C). This connection is further enhanced by hydrogen bonds involving Raf E174 and the backbone of Ras G48, and by direct interactions between Raf T178 and both Ras R149 on loop 10 and E153 on helix 5. Raf residues F163, F172, F141, and V180 create a hydrophobic patch in the core of the Raf-CRD that is in close contact with nonpolar residues in the central beta sheet of Ras, including V45, which also hydrogen-bonds to Raf S177 through its backbone amide. The Ras/Raf-CRD interaction is further strengthened by Q43, as its side chain extends toward the second Zn^2+^ binding site to make hydrogen bonds with both the backbone amide and carbonyl groups of H139 (Fig.1C). These key interactions involving Ras Q43 are complemented by hydrogen bonds donated by neighboring residue, K42, to the backbone carbonyl groups of both Raf V180, and T178 which in turn interacts directly with Ras N26 and, through a water molecule, with Ras I24 on helix 1. Ras Q25, positioned between I24 and N26 on helix 1 bridges the interacting surfaces of the two Raf domains by donating a hydrogen bond to the backbone carbonyl of Raf-RBD residue V88 and accepting one from the side chain of Raf-CRD T182. This binding mode of the Raf-CRD to Ras, shown clearly in our crystal structure, is distinct from that described in the recent NMR data-driven nanodisc-bound KRas/CRaf-RBD_CRD complexes (Fig. S1B), where the Ras/Raf-CRD interaction was modelled utilizing chemical shift perturbation data that are also consistent with our model (Fig. S1A) and paramagnetic relaxation enhancement (PRE) data with the unfortunate placement of the probe at a cysteine engineered in place of Q43 (*17*), an important residue at the Ras/Raf-CRD interface (Fig. 1C). In both structures, the Ras-binding interface on the Raf-CRD is separate from its proposed membrane-interacting regions (Fig. S1C). However, only the RBD-CRD configuration in our HRas/CRaf-RBD_CRD crystal structure is consistent with the expected location of the Raf-RBD when superimposed on the cryo-EM structure for the inactive BRaf/MEK1/14-3-3 complex (PDB ID 6NYB), in which the Raf-CRD is sandwiched between the Raf-kinase domain and 14-3-3 dimer and the Raf-RBD is solvent exposed, absent from the final cryo-EM reconstruction (Fig. S1D) (*27*).

### Raf-CRD links to the active site through loop 8 across the dimer interface

Given the importance of the Ras dimer in signaling through Raf, and the position of the CRD relative to helix 5 at the dimerization interface in the monomer of the HRas/CRaf-RBD_CRD complex, we modelled the HRas/CRaf-RBD_CRD dimer utilizing a two-fold symmetry axis conserved across a large collection of Ras crystal structures, including our structure of the HRas/CRaf-RBD complex (PDB ID 4G0N) (*7, 39, 40*). In the resulting model, the Raf-CRD is positioned for interaction with both Ras protomers in the Ras dimer, bridging the interswitch region of one to loop 8 of the other at the base of the Ras dimerization interface (Fig. 2A). A second dimer model was constructed utilizing a closely related helix 4/helix 5 Ras dimerization interface, recently described for NMR-data driven nanodisc-bound KRas dimers in the absence of Raf (PDB ID 6W4E) (Fig. S2A) (*41*). We performed three replicates of molecular dynamics simulations on both models. The model based on the crystal structures equilibrated quickly and was stable for all three simulations (350ns, 250ns, 250ns) (Fig. S2B, S2C), whereas the model based on the interface taken from the NMR-driven structure did not equilibrate within the first 75 ns, after which we stopped the simulations (Fig. S2B, S2D). We have previously shown robust, high-affinity dimerization of Ras in the presence of Raf (*38*), which may modulate the Ras/Ras interface in a manner that does not occur in the NMR model with data collected in the absence of Raf-RBD. Given these results, we proceeded with analysis of the simulations based on the model generated from the two-fold crystallographic symmetry taken from our structure of the Ras/Raf-RBD complex (PDB ID 4G0N). These simulations show greater fluctuations for Raf-CRD than observed for the rest of the complex (Fig. S2E), while it remains optimally positioned at the base of the Ras dimerization interface with potential to impact signaling and regulation of the complex.

**Fig. 2.**
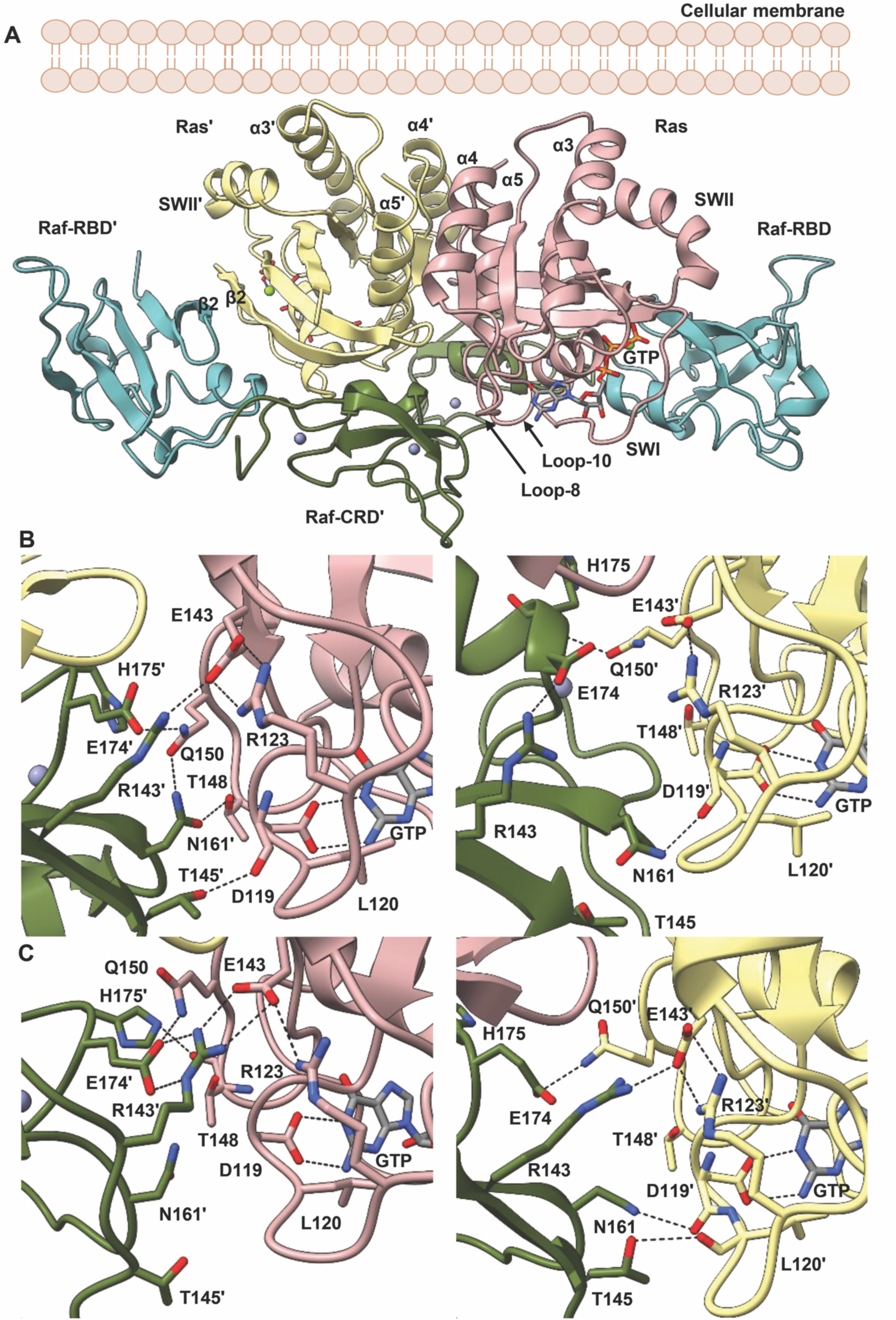
HRas/CRaf-RBD_CRD dimer complex. **(A)** Model dimer of the HRas/CRaf-RBD_CRD complex places the Raf-CRD at the base of the dimerization interface to interact with both Ras and Ras’ protomers. An orientation of the dimer with Ras helices 3, 4 and 5 roughly perpendicular to the membrane would exclude Raf-CRD insertion in the lipid bilayer. **(B-C)** Interactions at different time frames in the simulations illustrate the asymmetry introduced into the complex through the Raf-CRD’s dynamic interactions with the opposing Ras protomer. The left panels show Ras/Raf-CRD’ and the right panels show Ras’/Raf-CRD at representative time frames of (B) 60 ns into the replicate 2 simulations and (C) 140 ns into the replicate 3 simulations.

In the starting model constructed utilizing crystallographic symmetry, Raf-CRD residues R143, N161, and E174 are within interacting distance of loop 8 residue R123, the nucleotide binding residue D119, and Q150 at loop 10 immediately preceding helix 5 in the opposing Ras protomer. Ras D119 is part of the NKxD motif (residues 116-119), and directly coordinates the guanine base in the Ras active site (Fig. 2B, 2C). This motif is followed by loop 8 containing residue R123, which in turn makes a salt bridge interaction with E143 of the ExSAK motif (residues 143-147), linking two highly conserved regions important for active site stability (*8*). The Ras R123-E143 salt bridge remains present throughout the simulations, even as R143 of the Raf-CRD moves to interact with Ras E143 (Fig. 2B, 2C, Fig S3A (replicate 1), Fig. S4A (replicate 2) & Fig. S5A (replicate 3)), contributing favorably to the interface with the opposing Ras promoter. Raf-CRD residue N161 samples various states in which it can make contacts with the side chains of Ras T148 and Q150 in loop 10 (Fig. 2B, 2C, Fig. S3B, Fig. S4B & Fig. S5B) or with the backbone of loop 8 residues C118, D119, and L120 (Fig. 2B, 2C, Fig. S3C, Fig. S4C & Fig. S5C), while Ras Q150 also interacts with Raf-CRD E174 and H175 (Fig. 2B, 2C, Fig. S3D, Fig. S4D & Fig. S5D). Raf-CRD T145 is also observed to interact with the loop 8 residues D119 and L120 (Fig. 2B, 2C). The dynamic ability of the two Raf-CRDs to interact with the opposing Ras molecule provides a means to introduce some local asymmetry into the Ras/Raf dimer complex (Fig. 2B, 2C, Fig. S3A-D, Fig. S4A-D & Fig. S5A-D). There are several isoform-specific residues in this region of Ras (Fig. S6A, S6C) (*8*) and the flexibility of interaction between the Raf-CRD and loop 8 may be a key feature allowing the three Ras isoforms to have specific interactions with each of the Raf proteins (*42*). Furthermore, the asymmetry and flexibility that we observe in the dimer may facilitate a given Ras isoform to accommodate heterodimers of Raf, important in signal activation (*43*), as CRaf residues N161, E174, and H175 are isoform specific and correspond to Q257, Q270, and R271 in BRaf (Fig. S6B, S6C). Overall, in the Ras/Raf dimer, the Raf-CRD has access to the Ras dimerization interface through its interactions with loop 3 and helix 5, and to the Ras active site through loop 8, poised to affect key allosteric connections involved in the regulation of Ras (*44*).

### Allosteric communication across the Ras/Raf-RBD_CRD dimer complex

We recently demonstrated that the Raf-RBD promotes Ras dimerization and used dynamical network analysis and optimal/suboptimal path calculations based on MD trajectories to identify enhanced allosteric connections linking both Ras-dimerization and Raf-RBD binding interfaces in the HRas/CRaf-RBD dimer (*38*). By selecting residue D113 of each Raf-RBD for its long-range contributions on the Ras/Raf interaction (*45*), we observed a communication path extending 85 Å across the Ras/Raf-RBD dimer complex and identified residues D47, located in loop 3 of one Ras protomer, and E143, in β-strand 6 of the other, as a critical point of information transfer across the Ras dimerization interface (*38*). The significance of linking the two Raf-RBD residues D113 in the dimer of the Ras/Raf-RBD_CRD complex is suggested by its positioning, along with Raf-RBD D117, in the Raf-RBD binding pocket for the scaffold protein galectin that is critical for the activation of the MAPK pathway through a currently unknown mechanism (*46*).

It is intriguing that the Raf-CRD is located at the base of the Ras/Raf dimer to interact with both loop 3 of one Ras molecule and residue E143 of the other, therefore in contact with residues identified as critical for extending allosteric communication across the Ras dimerization interface (*38*). Due to longer simulation run times and the dynamic nature of the Ras/Raf-CRD interactions across the dimer interface (Fig. S3A-D, S4A-D & S5A-D), each trajectory was first divided into 50 ns segments prior to performing dynamical network analysis for the identification of residues with correlated motion within the dimer of the HRas/CRaf-RBD_CRD complex (*47, 48*). For each of the trajectory segments, each residue in the complex is represented by a spherical node positioned at the alpha carbon and edges are drawn between those whose atoms stay within 4.5 Å for over 75% of the MD trajectory (*49*). The allosteric connections revealed by the optimal/suboptimal paths calculated here in the presence of the Raf-CRD between the two D113 residues of each Raf-RBD in the dimer show four modes of intermolecular information transfer identified in our various trajectory segments that are consistent across the three replicates (Fig. 3A & S7A, S7B, S7C) (Table S2, S3 & S4). The first mode of information transfer involves the interswitch region and β-strand 6 as observed in the HRas/CRaf-RBD dimer simulations (*38*) (Fig. 3B, orange path). The second mode involves helix 5 and β-strand 6 as described for KRas/CRaf-RBD dimer simulations modelled on the membrane (*38*) (Fig. 3B, red path). The remaining two means for information transfer directly involve the Raf-CRD, with one utilizing the Raf-CRD’s close association with loop 8, creating paths that wrap around the Ras active site (Fig. 3C); and the other alternatively using the interconnectivity between the Raf-CRD and interswitch region to link the two Ras protomers again through loop 3 and β-strand 6 (Fig. 3D).

**Fig. 3.**
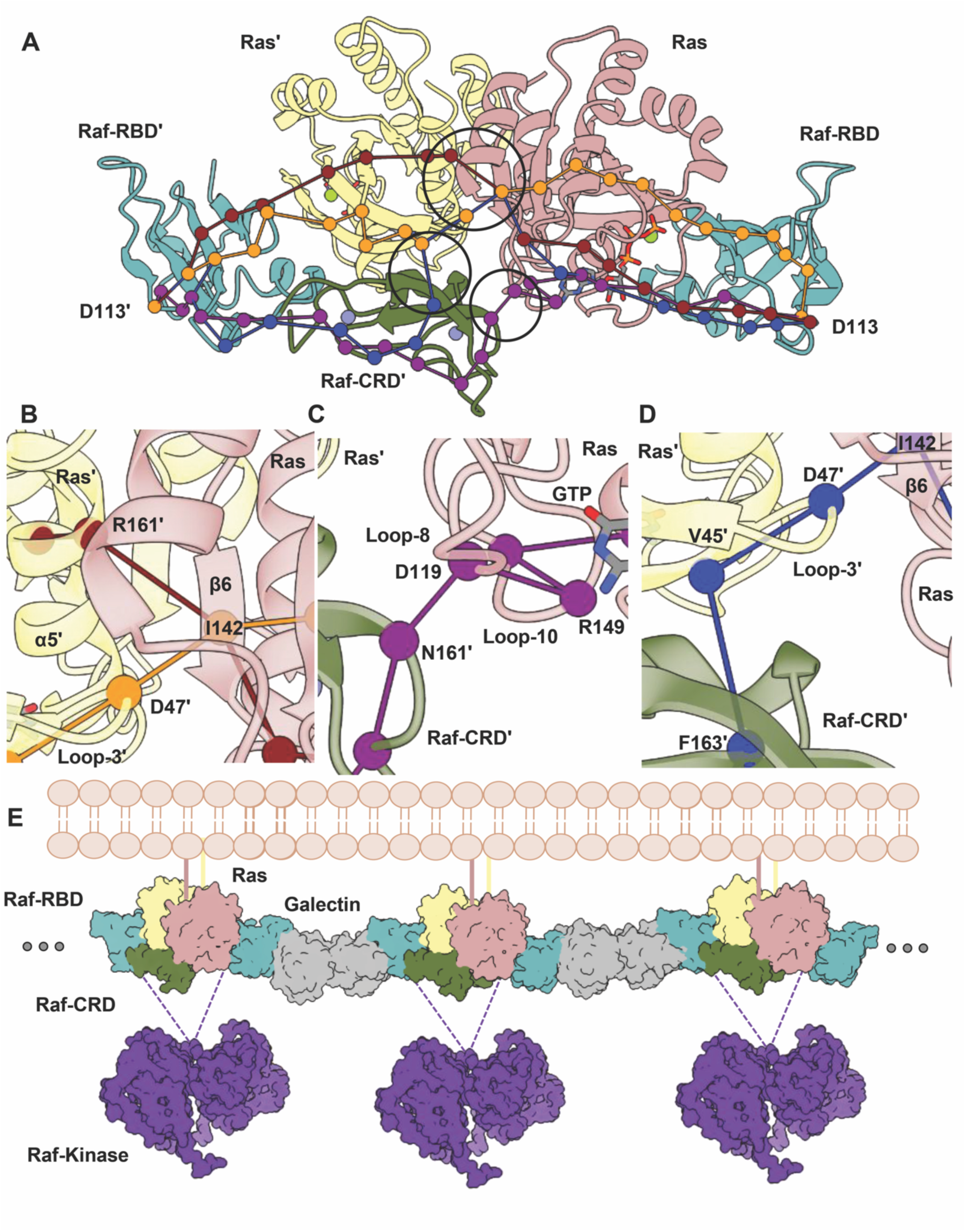
Dynamical network analysis performed for HRas/CRaf-RBD_CRD dimer simulations. **(A)** Optimal paths calculated between D113 and D113’ of each Raf-RBD molecule. **(B)** Close up view showing path for intermolecular information involving Ras loop 3/β-strand 6 (orange) and Ras helix 5/ β-strand 6 (red) across the Ras dimerization interface. **(C)** Close up view showing path for intermolecular information involving Raf-CRD/Ras loop 8 (purple) **(D)** Close up view showing path for intermolecular information involving Raf-CRD/Ras interswitch region (blue) **(E)** Proposed Ras/Raf/galectin assemblies adapted from Packer et al. (*38*) with inclusion of the Raf-CRD. Galectin (grey) (PDB ID 3W58) and Raf-kinase (purple) (PDB ID 6Q0J) were downloaded from the Protein Data Bank.

The allosteric connections linking residues D113 at the galectin binding pocket of each Raf-RBD molecule in the dimer, which also includes residue D117 (*46*) (Table S2, S3 & S4), leads to a model where Ras/Raf-RBD_CRD dimers couple with galectin dimers to form a higher order macromolecular platform involved in signal amplification and kinetic proofreading (Fig. 3E) (*38*), similar to that described for the LAT/Grb2/SOS system (*50*). This model is consistent with the functional importance of galectin for signaling through the MAPK pathway (*51-53*) and with results from single molecule tracking experiments correlating immobile species of Ras observed in live cells to active signaling through Ras/Raf/MEK/ERK (*54-56*).

Our molecular dynamics simulation data suggest that the interconnectivity between the Raf-CRD and the interswitch region of one Ras molecule and loop 8 of the other, optimally positions the Raf-CRD to diversify the paths of allosteric communication across the Ras/Raf complex linking the galectin binding sites at the two ends of the dimer, not only by further bridging the two Ras molecules in the Ras dimer, but also serving as a link between the Raf-RBD and second Ras molecule in the complex, with a direct connection to the Ras active site. Simulations of the KRas/CRaf-RBD dimer complex modelled on the membrane demonstrate that this complex is flexible at the interface (*38*). The Raf-CRD, poised to stabilize the base of the Ras dimerization interface and with involvement in intermolecular information transfer across the complex as identified in our optimal/suboptimal path calculations, may be a means to accommodate the complex’s dynamic nature by providing alternative routes to maintain communication between the two galectin-binding regions of each Raf-RBD, thereby establishing a platform most effective for signal transduction.

Interestingly, the location of the Raf-CRD in the context of the Ras/Raf dimer, combined with evidence supporting an approximately perpendicular orientation of helices 3, 4 and 5 with respect to the membrane (*38, 39*), precludes insertion of the Raf-CRD into the membrane. In light of our Ras/Raf-RBD_CRD structure and the increasing evidence that the helix 4/helix 5 Ras dimer is an essential unit for signaling through Ras/Raf/MEK/ERK (*34, 38-41*), we must question the presumed membrane-binding function of the Raf-CRD that has been characterized only in the context of monomeric Ras/Raf-RBD_CRD complexes (*17, 18, 20*). Instead we suggest that the primary function of previously proposed hydrophobic membrane-binding region, conserved across Raf isoforms, is to stabilize Raf autoinhibition in the absence of Ras as demonstrated in the inactive BRaf/MEK1/14-3-3 cryo-EM structure (*27*).

### Mechanism for Ras-mediated release of Raf autoinhibition

The cryo-EM structure of the inactive BRaf/MEK1/14-3-3 complex (PDB ID 6NYB) demonstrates that the autoinhibited Raf state is maintained through the interaction of the 14-3-3 dimer with phosphoserine residues located in Raf CR2 and CR3, and the contacts established between the centrally located Raf-CRD with both the Raf kinase domain and 14-3-3 dimer (*27*). The 14-3-3 and kinase-domain interfaces on the Raf-CRD involve hydrophobic and charged residues that are proposed to interact with the membrane (*27*) and do not overlap with the Ras-binding region revealed in our crystal structure (Fig. 4A). The active BRaf/MEK1/14-3-3 structure (PDB ID 6Q0J), in the absence of the CR2 14-3-3 binding motif, displays a shift of the 14-3-3 dimer with respect to the Raf-kinase domain. This shift is accompanied by conformational rearrangement of the Raf-kinase C-terminus, which now redirects away from the Ras-binding surface of the Raf-CRD (Fig. 4B).

**Fig 4.**
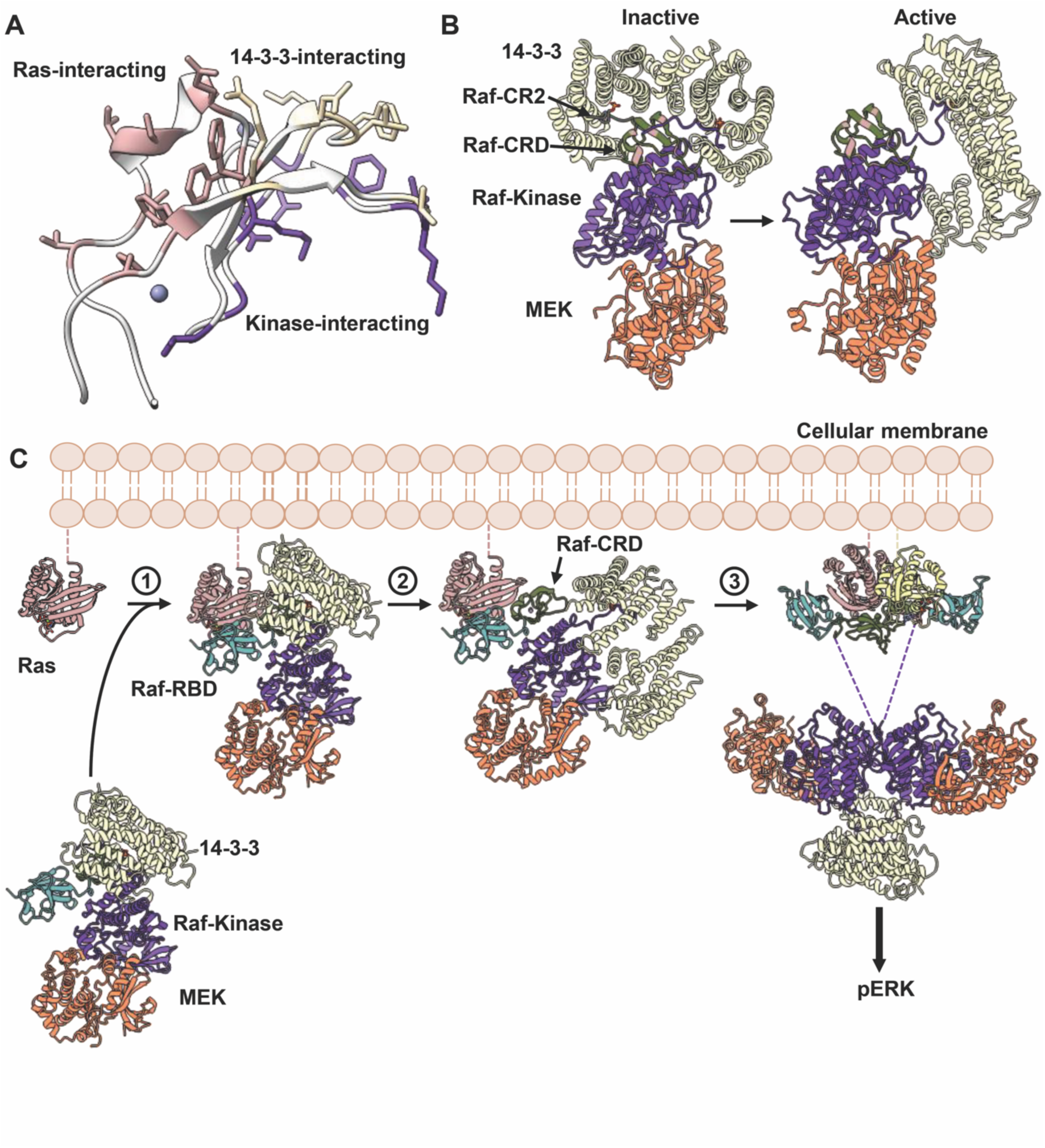
Proposed Mechanism for Ras-mediated release of Raf autoinhibition. **(A)** Residues involved in the Ras/Raf-CRD interaction (pink) do not overlap with Raf-CRD residues used to stabilize contacts with the kinase domain (purple) and 14-3-3 proteins (wheat) in the autoinhibited state. **(B)** Comparison of Raf-kinase domain and 14-3-3 conformations in the inactive (6NYB) and active Raf structure (6Q0J). Note the shift in the 14-3-3 dimer in the absence of the Raf CR2 motif and release of the Raf-CRD **(C)** Proposed structural model for the Ras-mediated release of Raf autoinhibition for MAPK signaling. Raf-RBD is exposed in the autoinhibited state. Its disorder in the cryo-EM structure is likely due to delocalization provided by the linker to the CRD. The high affinity binding of Raf-RBD to Ras places the exposed area of the CRD in its Ras binding site, while steric hindrance is expected to displace the Raf C-terminus and promote the shift of the 14-3-3 dimer towards its active state interaction with a second Raf kinase domain. These two features of the Ras/Raf interaction are shown in steps 1 and 2. Binding of Raf-RBD concurrently promotes dimerization of the complex as shown in step 3, which we propose is a key step in binding galectin dimers to build a platform for signal amplification as shown in Fig. 3E.

The absence of the Raf-RBD from the inactive BRaf cryo-EM reconstruction suggests it does not contribute toward stabilizing the autoinhibited state and therefore must remain accessible for Ras-binding. Indeed, by aligning the CRaf-CRD from our Ras/Raf-RBD_CRD structure with the BRaf-CRD in the inactive complex, we observe that the short Raf-RBD_CRD linker is sufficient for the Raf-RBD to be accessible for interaction with Ras (Fig. 4C & Fig. S1D).

It has previously been shown that the affinity of 14-3-3 proteins for the CR2 site is weaker than that for the CR3 binding motif (*24*). We propose that Ras-binding to the Raf-RBD induces conformational rearrangements of both Raf CR2 and CR3 regions that result in the dissociation of the 14-3-3 subunit from the CR2 binding site. The redirection of the Raf C-terminus may be coupled to the dissociation of 14-3-3 from the Raf CR2 binding motif, facilitated by the interaction with Ras. This would expose the CR2 site for dephosphorylation, preventing association of the 14-3-3 dimer to reestablish Raf autoinhibition. This is consistent with an increase in Raf-kinase activity in cells upon dephosphorylation of the CR2 site (*23, 24, 28*). The displacement of 14-3-3 from the CR2 binding site would promote its interaction with a second Raf kinase to facilitate kinase domain dimerization and further expose the Raf-CRD for extraction (Fig. 4B, 4C), thus resulting in complete collapse of the autoinhibited conformation and progression toward Raf activation.

In our proposed model for Ras-mediated activation of the MAPK pathway (Fig. 4C), binding of Raf-RBD to Ras promotes Ras dimerization to facilitate galectin binding through allosteric modulation of its binding site on Raf-RBD, resulting in a platform of Ras/Raf and galectin dimers for signal amplification (Fig. 3E). During this process the CRD is released from its autoinhibitory role, allowing the shift in 14-3-3 and stabilization of the Raf dimer. Given the high-affinity dimerization of Ras promoted by Raf-RBD that we previously reported (*38*), we propose that these events occur concertedly, resulting in simultaneous release of autoinhibition and allosteric modulation of the galectin binding site on Raf-RBD, with robust activation of the Ras/Raf/MEK/ERK pathway.

## Conclusions

The crystal structure of the Ras/Raf-RBD_CRD complex presented here provides a key missing link in our understanding of Ras-mediated Raf activation. It shows a continuous surface for Ras/Raf interaction that links the Ras active site and dimerization interface through switch I, loop 3, and helix 5. In the context of the dimer, the CRD also comes in contact with loop 8 and residues in two highly conserved nucleotide-binding motifs, centrally positioned to access regulatory and functional regions of Ras. This is consistent with the observation that the Raf-CRD is required for Raf activation *in vivo*. Our analysis of this structure in the context of other recent structural breakthroughs in the literature, provides an emerging model of Ras activation of Raf where formation of the Ras/Raf complex is linked to both dimerization and release of Raf autoinhibition to form a platform for effective signaling. Our structure and analysis put in question the presumed membrane-binding function of the Raf-CRD, at least in the context of the dimer required for signaling through Raf. Instead, we build upon our recent proposal of higher order Ras/Raf/galectin signaling assemblies, paving forward a new direction for future investigations that will be paramount to our understanding of signaling through Ras/Raf/MEK/ERK. Because the affinity between Ras and the Raf-CRD alone is relatively weak, the disruption of its interactions with Ras, either through the interswitch region involving loop 3 or through loop 8, holds strong potential as a novel approach for targeting Ras and Raf related cancers. The structural information provided here, with increased mechanistic understanding of Raf activation, provides a new framework from which to explore the effects of oncogenic mutants on signaling through the MAPK pathway.

## Acknowledgments

We thank Alicia Volmar for performing site directed mutagenesis on WT HRas to obtain the HRasR97C mutant used to crystallize the complex with Raf-RBD_CRD. We thank Kathleen Merritt for helping to make a series of constructs of the Raf-RBD_CRD used in crystallization trials to obtain the structure presented here. We are grateful to all members of the Mattos lab for helpful discussion and sharing of computational resources. We acknowledge Northeastern University Research Computing for our usage of the Discovery cluster to perform all molecular dynamics simulations described and analyzed in this paper.

## Funding

This work was funded by the National Science Foundation MCB-1517295 grant awarded to C.M. and the 2018 Fall Research and Creative Endeavor Award and 2019 Summer Scholars Independent Research Fellowship awarded to T.C. by the Northeastern University Office of Undergraduate Research and Fellowships.

## Author contributions

T.C. made protein constructs, purified protein, collected X-ray data, performed structure refinement and analysis, performed molecular dynamics simulations and analysis, and wrote the paper. C.M. conceived the project, performed structure analysis, and wrote the paper.

## Competing interests

Authors declare no competing interests.

## Data and materials availability

Crystal structure coordinates and structure factors are available from the PDB under accession code 7JHP.

## Materials and Methods

### Protein Expression/Purification

The HRas/CRaf-RBD-CRD complex was crystallized using the HRasR97C allosteric site mutant. The HRasR97C construct containing residues 1-166 was generated by PCR site-directed mutagenesis from the wild type HRas construct similarly truncated. HRas protein was expressed, purified, and GDP was exchanged for the non-hydrolyzable GTP analog GppNHp as previously described (*57*). The GB1_Raf-RBD-CRD (52-184) construct as previously published (*7, 58*) was extended utilizing PCR site-directed mutagenesis to generate the construct of CRaf containing residues 52-187 used for the experiments described in this paper. CRaf protein was transformed into *Escherichia coli* BL21(DE3) cells and grown at 37°C at 220 rpm in the presence of 50 mg/L ampicillin. Expression was induced in the presence of 20 μM ZnCl2 at an OD_600_ of 0.6-0.8 with 0.5 mM IPTG. The temperature was lowered to 32°C and cells were harvested after 5 hours by centrifugation.

Cells were resuspended in 50 mM Tris-HCl pH 7.4, 500 mM NaCl, 1 mM BME, 5% v/v glycerol, and 20 μM ZnCl2 in the presence of 1 mg/mL leupeptin, 1 mg/mL pepsatin, and 12 mg bezamidine. Cells were lysed by sonication and the insoluble fraction was separated by centrifugation at 13,500 rpm, 4°C for 30 minutes. The supernatant was syringe filtered through a 0.45 μm membrane before performing Ni NTA affinity chromatography (HisTrap HP, GE Lifesciences). CRaf protein was eluted with 200 mM imidazole. Fractions were pooled in the presence of 1 molar equivalent HRasR97C (1-166) and concentrated to 2 mL. Protein was dialyzed overnight into cleavage buffer (20 mM Tris-HCl pH 8.1, 100 mM NaCl, 1 mM DTT, 25 mM CaCl_2_, 5 mM MgCl_2_, 20 μM ZnCl_2_, 5% v/v glycerol) at 4°C. 10 units of thrombin were added per mg of CRaf_52-187 and incubated for 18 hours at room temperature. The cleaved complex was further purified by size exclusion chromatography (16/60 Sephacryl S100, GE Lifesciences).

### Protein Crystallization, Data Collection, Refinement

The HRasR97C_1-166/CRaf_52-187 complex was concentrated to 7 mg/ml for crystallization trials. Crystals of the complex were obtained by sitting-drop vapor diffusion with reservoir solution of 0.1 M ammonium acetate, 0.1 M Bis-Tris pH 5.5, and 17% w/v PEG 10K. Drops were prepared with 1 μL protein and 1 μL reservoir solution and crystals were grown at 18°C over ten weeks. Data collection was performed on a home source MicroMax007HF with a Cu^2+^ anode, tungsten filament, and R-AxisIV^2+^ detector from Rigaku. Data was indexed, integrated, and scaled with the HKL3000 software package *(58)*. Molecular replacement was performed in PHENIX (*59*) utilizing the HRas/CRaf-RBD crystal structure (PDB ID 4G0N) as the search model. After correct placement of Ras and Raf-RBD molecules, an additional search was performed for the Raf-CRD utilizing the Raf-CRD NMR structure (PDB ID 1FAR). Further structure refinement was performed with PHENIX (*59*) and COOT (*60*).

### Model Preparation for Molecular Dynamics Simulations

The monomer of the HRas/CRaf-RBD_CRD complex was prepared by replacing the fully ordered HRas G-domain (PDB ID 3K8Y) for the Ras G-domain in the HRasR97C/CRaf-RBD_CRD crystal structure. Loop 4 of the Raf-RBD (residues 103-109) was modelled using the coordinates of the Raps/Raf-RBD crystal structure (PDB ID 1C1Y). The Raf-RBD_CRD linker region and unmodelled side chains were built using guidance from the 2Fo-Fc electron density map contoured at lower sigma levels.

To obtain the dimer model with the helix 4 and helix 5 interface frequently observed in crystal structures, the dimer generated by 2-fold crystallographic symmetry from our structure of HRas/CRaf-RBD (PDB ID 4G0N) was used as a template for superposition of two HRas/Raf-RBD_CRD complexes to generate the dimer containing the CRD. A second dimer model was generated by aligning the Ras/Raf-RBD_CRD monomer model with the coordinates of each Ras G-domain in the NMR-driven model of KRas-GTP tethered to a lipidated nanodisc (PDB ID 6W4E).

The six cysteines located in the CRaf-CRD for all systems were patched with CYN for stable coordination of the two Raf-CRD zinc ions throughout the simulations. All histidines in both HRas and CRaf molecules were in the uncharged HSE form.

### Molecular Dynamics Simulations

Each system was solvated in a TIP3P water box and sodium and chloride ions were added to a final concentration of 150 mM to neutralize the system. Each system was minimized for 5000 steps and gradually heated from 50 to 250 K prior to the production runs. Production runs were performed at constant temperature and pressure of 300 K and 1.01325 bar with periodic boundary conditions. The first 30 ns of each simulation had a time step of 1 fs and the remaining simulation time was performed with a time step of 2 fs. Simulations were performed with the NAMD package (*61*) and CHARMM27 force field (*62*). The Particle Mesh Ewald method with a grid size of 1 Å was used to calculate long-range electrostatics. Simulations of each system were performed in triplicate. Simulations for the crystallographic dimer model are denoted as D1, D2, and D3 throughout the text for 350, 250, and 250 ns. Simulations for the NMR data-driven dimer model are referred to as N1, N2, N3 for a total length of 75 ns, after which the simulations failed to equilibrate and were terminated.

### Trajectory Analysis

Dynamical network analysis was performed using the Carma (*63*) and Catdcd software packages. Each protein residue within the simulation is assigned a spherical node centered on its alpha carbon. Edges are drawn between nodes that stay within 4.5 Å for at least 75% of the trajectory and are weighted utilizing pairwise cross correlation data. Optimal and suboptimal path calculations can also be used to identify residues important in allosteric communication. The optimal path is calculated as the path containing the fewest number of edges connecting user specified “source” and “sink” nodes. Nodes occurring most frequently in the calculated optimal and suboptimal paths are important for allosteric communication between the two specified “source” and “sink” nodes (*49*).

Root-mean square deviation (RMSD) and root-mean square fluctuation (RMSF) calculations were performed using the Bio3d software package (*64*). Distance calculations were performed in VMD (*65*) by sourcing the distance.tcl script that can be found in the VMD script library.

## Notes

### Competing Interest Statement

The authors have declared no competing interest.

