## Supplemental Tables and Figures for "Crystal structure reveals the full Ras:Raf interface and advances mechanistic understanding of Raf activation"

### Supplementary Information

**This PDF file includes**

Tables S1 to S4
Figures S1 to S7

**Table S1. X-Ray data collection and structure refinement statistics**

| HRas/CRaf-RBD_CRD complex (PDB 7JHP) | |
|---|---|
| **Data Collection** | |
| Space group | C 1 2 1 |
| Unit cell dimensions | |
| *a, b, c* (Å) | 94.49, 44.35, 74.40 |
| α, β, γ (˚) | 90, 103.95, 90 |
| Resolution (Å) | 36.61-2.77 (2.87-2.77) |
| I/$\sigma$ | 15.63 (2.08) |
| Completeness (%) | 91.48 (49.27) |
| Redundancy | 7.2 (6.9) |
| $R_{merge}$ | 0.120 (0.785) |
| **Refinement** | |
| No. reflections | 349414 |
| Unique reflections | 7158 (369) |
| $R_{work}$ | 0.21 (0.26) |
| $R_{free}$ | 0.26 (0.34) |
| No. atoms | 2311 |
| Macromolecules | 2228 |
| Ligands | 35 |
| Solvent | 48 |
| RMS(bonds) | 0.002 |
| RMS(angles) | 0.45 |
| Ramachandran (%) | |
| favored | 96.48 |
| allowed | 3.17 |
| outliers | 0.35 |
| Average B-factor | 50.88 |
| Macromolecules | 51.25 |
| Ligands | 40.16 |
| Solvent | 41.39 |

*Values in parentheses are for highest-resolution shell.

**Table S2. Residues involved in intermolecular information transfer at the Ras dimerization interface identified in optimal/suboptimal path calculations between Raf-RBD residues D113-D113' and D117-D117' for replicate 1.**

| 1 | D113-D113' paths | D117-D117' paths |
|---|---|---|
| **0-50 ns** | 465 paths<br>Ras G48-Ras' S127 80.4%<br>Raf N161 – Ras' D119/L120 13.9%<br>Ras D47-Ras' I142 2.9%<br>Raf S177-Ras V45 2.4% | 652 paths<br>Ras D47/G48-Ras' S127 92.3%<br>Ras D47-Ras' I142/E143 6.9%<br>Ras R161-Ras' I142 0.3% |
| **50-100 ns** | 864 paths<br>Raf L160/N161-Ras' D119 60.7%<br>Ras D47/G48-Ras' I142/E143 35.4% | 116 paths<br>Ras D47/G48-Ras' I142/E143 89.6%<br>Ras G48-Ras' S127 8.6% |
| **100-150 ns** | 2209 paths<br>Raf E174 – Ras' Q150 100% | 8369 paths<br>Raf E174 – Ras' Q150 100% |
| **150-200 ns** | 1902 paths<br>Ras C118 - Raf' N161 72.4%<br>Ras D119 - Raf' N161 24.6% | 1438 paths<br>Ras C118 - Raf' N161 71.3%<br>Ras D119 - Raf' N161 28.7% |
| **200-250 ns** | 1923 paths<br>Ras C118/D119 - Raf' L160/N161 58.2%<br>Ras C118/D119 - Raf' T145/F146 35.7%<br>Ras T148 - Raf' L160 6.1% | 6878 paths<br>Ras C118/D119 - Raf' L160/N161 60.4%<br>Ras C118/D119 - Raf' T145/F146 34.1%<br>Ras T148 - Raf' L160 5.4% |
| **250-300 ns** | 1923 paths<br>Ras D47 - Ras' I142/E143 100% | 6878 paths<br>Ras D47 - Ras' I142/E143 100% |
| **300-350 ns** | 265 paths<br>Ras R161 - Ras' I142 68.3%<br>Ras T148 - Raf' N161 16.6%<br>Ras D47/G48 - Ras' I142/E143 15.1% | 78 paths<br>Ras R161 - Ras' I142 75.6%<br>Ras D47/G48 - Ras' I142/E143 23.4% |

*Percentages were calculated as the number of times each edge appeared in the suboptimal path calculations divided by the total number of calculated paths.

**Table S3. Residues involved in intermolecular information transfer at the Ras dimerization interface identified in optimal/suboptimal path calculations between Raf-RBD residues D113-D113' and D117-D117' for replicate 2.**

| 2 | D113-D113' paths | D117-D117' paths |
|---|---|---|
| **0-50ns** | 1871 paths<br>Ras D47-Ras' I142/E143 100% | 1195 paths<br>Ras D47-Ras' I142/E143 99.4% |
| **50-100 ns** | 313 paths<br>Ras D47-Ras' I142 87.5%<br>Ras R161-Ras' I142 12.5% | 1110 paths<br>Ras D47-Ras' I142/E143 92.2%<br>Ras R161-Ras' I142 7.8% |
| **100-150 ns** | 155 paths<br>Ras C118-Raf' N161 100% | 111 paths<br>Ras C118-Raf' N161 100% |
| **150-200 ns** | 9897 paths<br>Ras R161-Ras' I142 90.6%<br>Ras D47-Raf' I142/E143 8.5%<br>Raf N161-Ras' 121 0.8% | 2099 paths<br>Ras R161-Ras' I142 97.0%<br>Ras D47-Raf' I142/E143 2.4% |
| **200-250 ns** | 78 paths<br>Raf T178-Ras V45 93.5%<br>Ras D47-Ras' I142 100% | 96 paths<br>Raf T178-Ras V45 94.7%<br>Ras D47-Ras' I142 99.0% |

*Percentages were calculated as the number of times each edge appeared in the suboptimal path calculations divided by the total number of calculated paths.

**Table S4. Residues involved in intermolecular information transfer at the Ras dimerization interface identified in optimal/suboptimal path calculations between Raf-RBD residues D113-D113' and D117-D117' for replicate 3.**

| 3 | D113-D113' paths | D117-D117' paths |
|---|---|---|
| **0-50ns** | 87 paths<br>Ras D154-Ras' D154 97.7% | 230 paths<br>Ras I142/E143-Ras' D47 55.6%<br>Ras D154-Ras' D154 42.4% |
| **50-100 ns** | 1083 paths<br>Raf E174-Ras V45 45.8%<br>Raf S177-Ras V45 36.6%<br>Raf F163-Ras V45 17.6%<br>Ras D47-Ras' I142 100% | 1839 paths<br>Raf E174-Ras V45 42.9%<br>Raf S177-Ras V45 38.8%<br>Raf F163-Ras V45 14.7%<br>Ras D47-Ras' I142 99.9% |
| **100-150 ns** | 420 paths<br>Ras R161-Ras' I142 78.3%<br>Ras D47-Raf' I142/E143 21.2% | 147 paths<br>Ras R161-Ras' I142 100% |
| **150-200 ns** | 8692 paths<br>Ras L120-Raf' R143 87.0%<br>Ras D119-Raf' R143 10.8%<br>Ras C118-Raf' R143 1.5%<br>Ras A121-Raf'143 0.6% | 10260 paths<br>Ras L120-Raf' R143 84.0%<br>Ras D119-Raf' R143 12.3%<br>Ras C118-Raf' R143 3.0%<br>Ras A121-Raf'143 0.7% |
| **200-250 ns** | 151 paths<br>Ras I142/E143-Ras' D47 100% | 695 paths<br>Ras I142-Ras' R161 56.7%<br>Ras I142-Ras' D47 43.2% |

*Percentages were calculated as the number of times each edge appeared in the suboptimal path calculations divided by the total number of calculated paths.

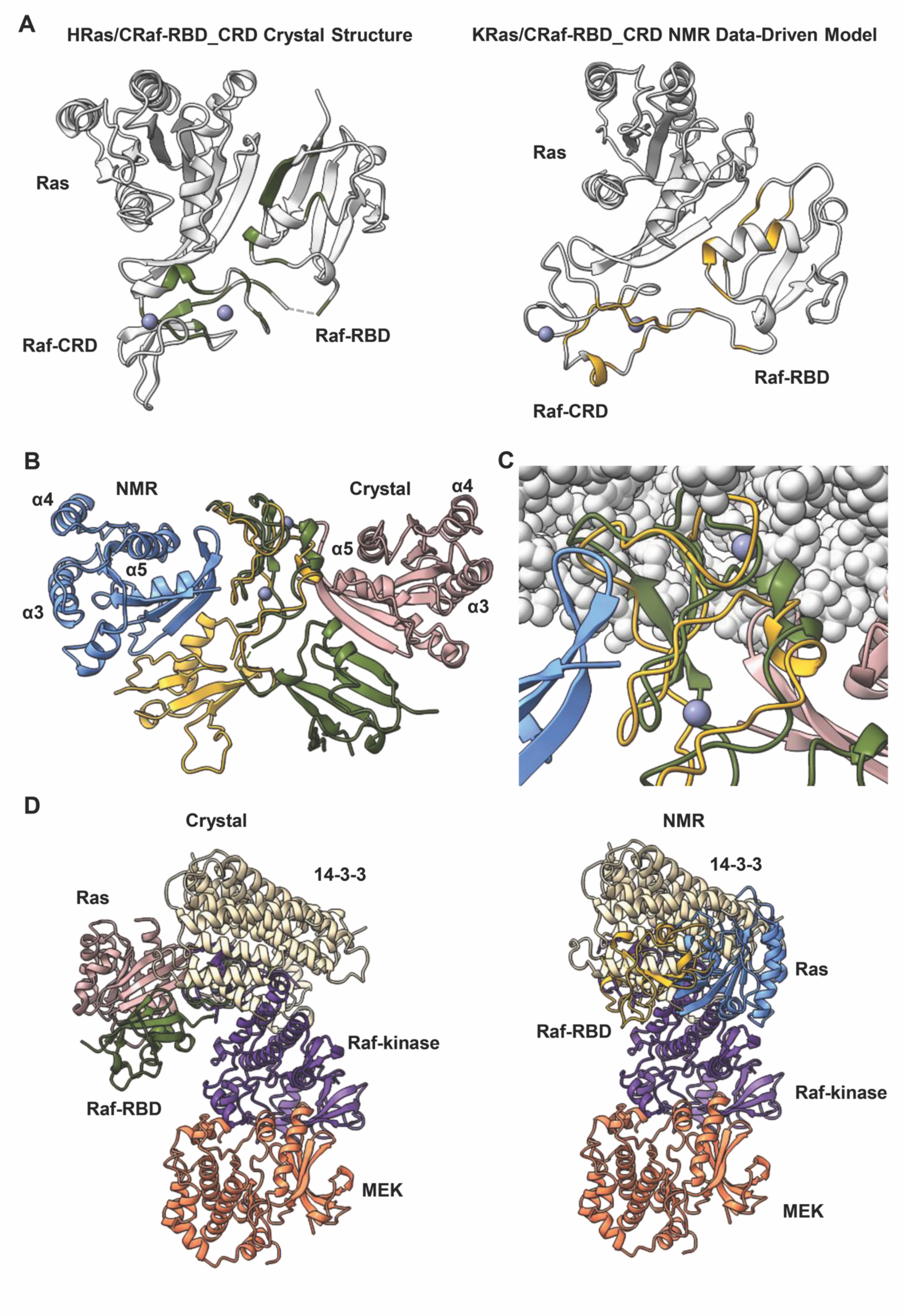
A
HRas/CRaf-RBD_CRD Crystal Structure
KRas/CRaf-RBD_CRD NMR Data-Driven Model
Ras
Raf-CRD
Raf-RBD
Ras
Raf-RBD
Raf-CRD
B
NMR
Crystal
α4
α5
α3
α4
α5
α3
C
D
Crystal
NMR
14-3-3
Ras
Raf-kinase
Raf-RBD
MEK
14-3-3
Ras
Raf-RBD
Raf-kinase
MEK

**Fig. S1**. **Comparison of the HRas/CRaf-RBD_CRD crystal structure with nanodisc-bound KRas/CRaf-RBD_CRD NMR data-driven structure (A)** Chemical shift perturbations used to model the nanodisc-bound KRas/CRaf-RBD_CRD structure (PDB ID 6PTS) mapped onto our HRas/CRaf-RBD_CRD crystal structure (left, green) and the KRas/CRaf-RBD_CRD NMR-driven model (right, yellow). Our crystal structure is consistent with the NMR data used to determine the KRas/CRaf-RBD_CRD model. **(B)** Alignment of the Raf-CRDs from our crystal structure (Ras, pink/Raf-RBD green) with that of the NMR model (Ras blue/Raf-RBD yellow) shows that the Raf-CRD is rotated 180˚ relative to Ras. **(C)** Close up view of the nanodisc/Raf-CRD interaction demonstrates that the Raf-CRD region interacting with the nanodisc membrane in the KRas/CRaf-RBD_CRD NMR model is still available for interaction with the membrane in our HRas/CRaf-RBD_CRD crystal structure in the absence of Ras dimerization. **(D)** Comparison of our HRas/CRaf-RBD_CRD crystal structure (left) and the KRas/CRaf-RBD_CRD NMR model (right) aligned with the BRaf-CRD from the inactive BRaf/14-3-3/MEK1 complex (PDB ID 6NYB). Only the Ras-binding interface identified in our crystal structure is accessible for Ras interaction in the autoinhibited state. Alignment of the Raf-CRDs between the NMR model and autoinhibited structure results in the placement of the Raf-RBD in a position inconsistent with the cryo-EM data *(26)* in which the Raf-RBD was solvent exposed and disordered.

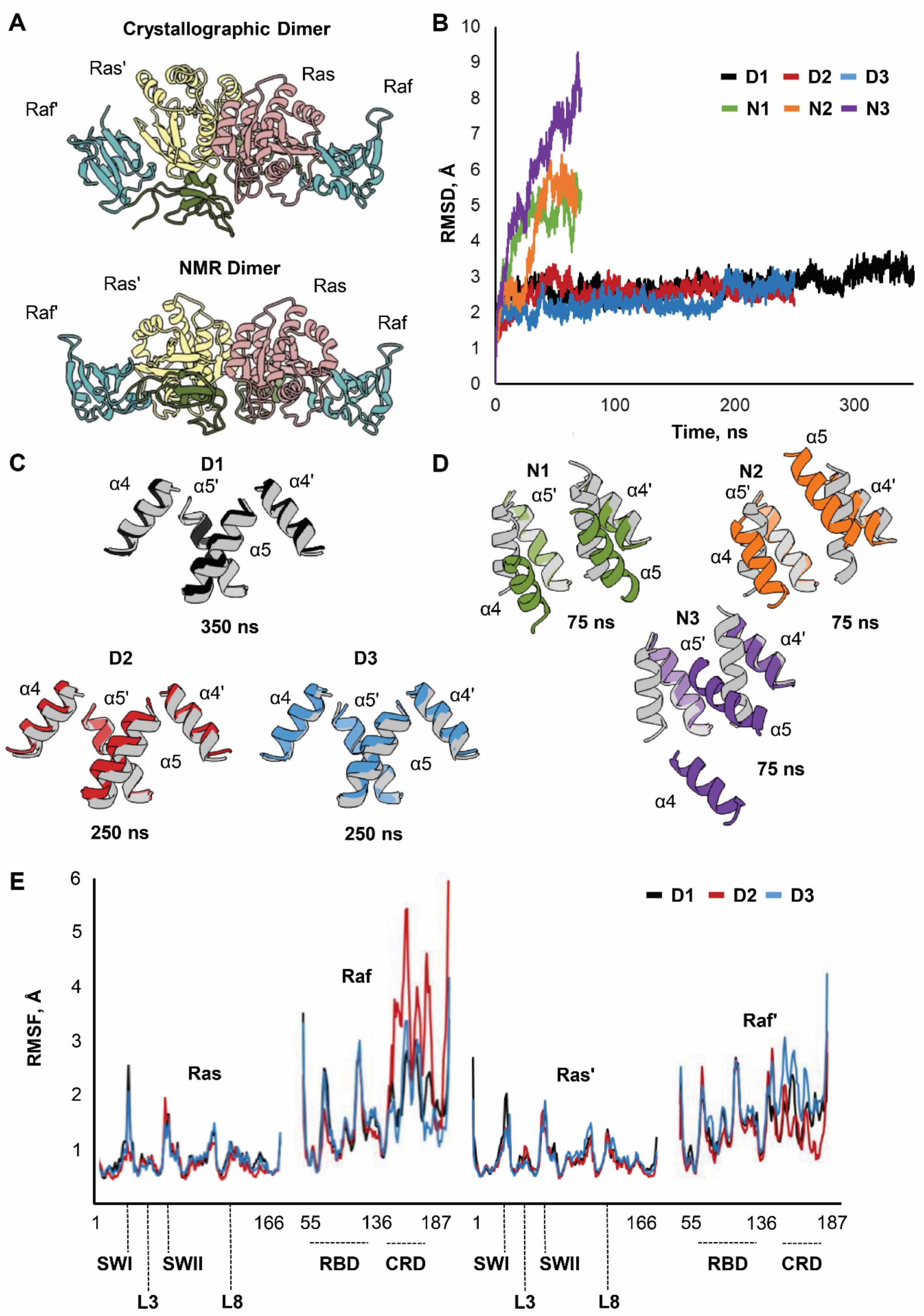
A
Crystallographic Dimer
Ras'
Ras
Raf'
Raf
NMR Dimer
Ras'
Ras
Raf'
Raf
B
RMSD, Å
Time, ns
D1 D2 D3
N1 N2 N3
C
D1
α4
α5'
α4'
α5
350 ns
D2
250 ns
D3
250 ns
D
N1
75 ns
N2
75 ns
N3
75 ns
E
RMSF, Å
D1 D2 D3
Ras
Raf
Ras'
Raf'
SWI
SWII
L3
L8
RBD
CRD

**Fig. S2. Comparison of MD simulations starting from crystallographic and NMR data driven helix 4/helix 5 Ras dimerization interfaces (A)** Starting models constructed for the HRas/CRaf-RBD_CRD dimer utilizing 2-fold symmetry in the HRaf/CRaf-RBD crystal structure (PDB ID 4G0N) (top) and nanodisc-bound KRas-GTP NMR data-driven dimer structure (PDB ID 6PTS) (bottom). **(B)** Root-mean square deviation (RMSD) shows 3 replicates for simulations of the dimer modelled with crystallographic symmetry (D1, D2, D3) that converge and are stable throughout the simulation time (350, 250, 250 ns). RMSD of simulations starting from the NMR dimer orientation (N1, N2, N3) do not equilibrate within 75 ns of simulation time. **(C)** Alignment of crystallographic symmetry derived dimer starting structure (grey) with the final frame from each trajectory shows that the dimer orientation remains stable throughout the simulation time. **(D)** Alignment of NMR-dimer starting structure (grey) with the final frame from each trajectory highlights the instability of this Ras/Raf-RBD_CRD dimer orientation. **(E)** Root-mean square fluctuation (RMSF) analysis for the crystallographic symmetry derived dimer simulation replicates (D1, D2, D3) shows enhanced fluctuations in the Raf-CRD that vary among replicates consistent with the dynamic variability of Ras/Raf-CRD interactions at loop 3 and loop 8 involving both Ras protomers.

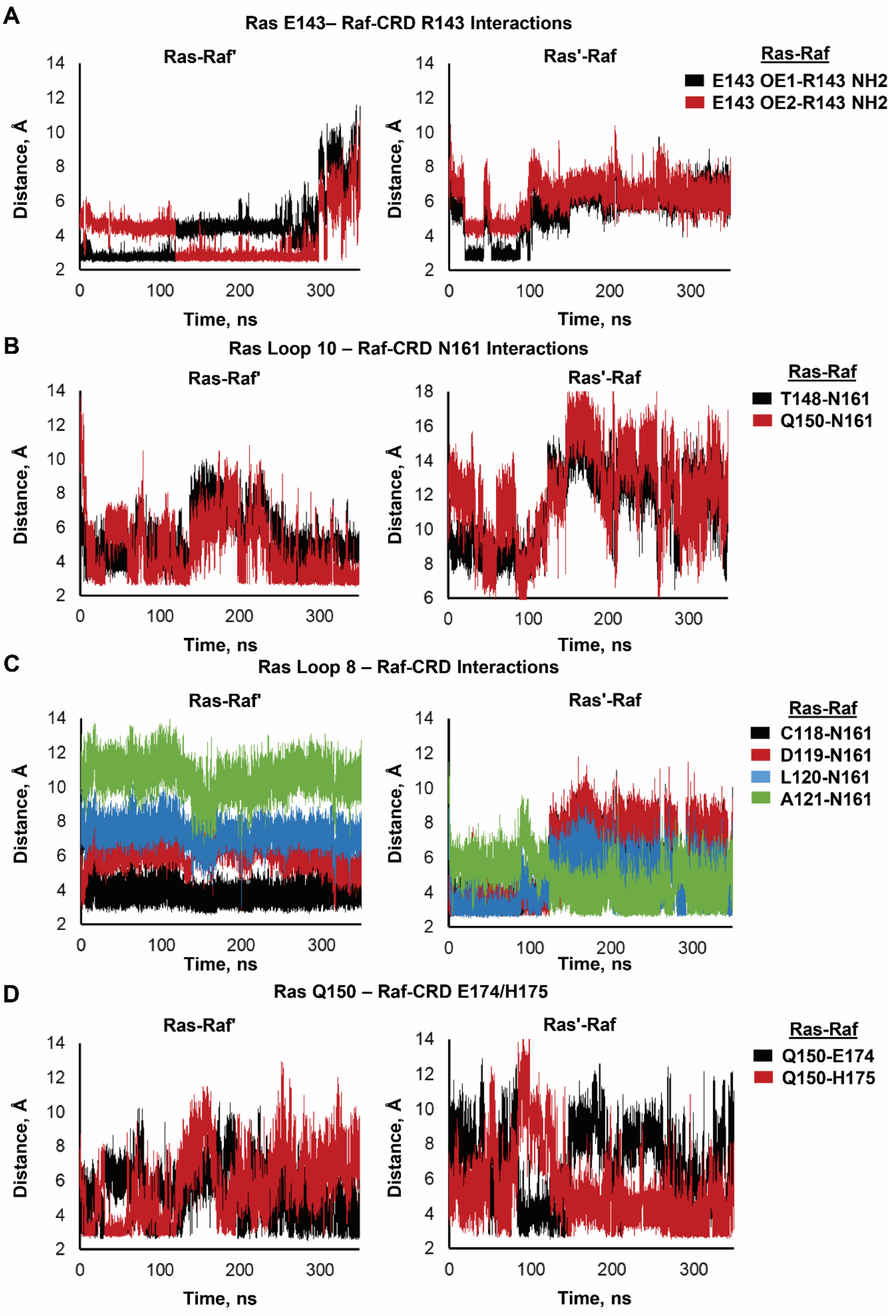
A
Ras E143– Raf-CRD R143 Interactions
Ras-Raf'
Ras'-Raf
Ras-Raf
E143 OE1-R143 NH2
E143 OE2-R143 NH2
Distance, Å
Time, ns
B
Ras Loop 10 – Raf-CRD N161 Interactions
Ras-Raf'
Ras'-Raf
Ras-Raf
T148-N161
Q150-N161
Distance, Å
Time, ns
C
Ras Loop 8 – Raf-CRD Interactions
Ras-Raf'
Ras'-Raf
Ras-Raf
C118-N161
D119-N161
L120-N161
A121-N161
Distance, Å
Time, ns
D
Ras Q150 – Raf-CRD E174/H175
Ras-Raf'
Ras'-Raf
Ras-Raf
Q150-E174
Q150-H175
Distance, Å
Time, ns

**Fig. S3. Ras/Raf-CRD' and Ras'/Raf-CRD interactions over 350 ns simulation time of the HRas/CRaf-RBD_CRD dimer complex (replicate 1). (A)** Distances calculated between the sidechain of Raf-CRD residue R143 and the side chain of residue E143 of the opposing Ras protomer. Ras/Raf-CRD' interactions are shown on the left and Ras'/Raf-CRD interactions are shown on the right. **(B)** Distances calculated between the sidechain of Raf-CRD residue N161 and the side chains of T148 (black) and Q150 (red) located in loop 10 of the opposing Ras protomer. **(C)** Distances calculated between the sidechain of Raf-CRD residue N161 and the carbonyl backbone of residues C118 (black), D119 (red), L120 (blue), and A121 (green) in the opposing Ras molecule. **(D)** Distances calculated between the sidechains of Raf-CRD residues E174 (black) and H175 (red) and the side chain of residue Q150 of the opposing Ras protomer.

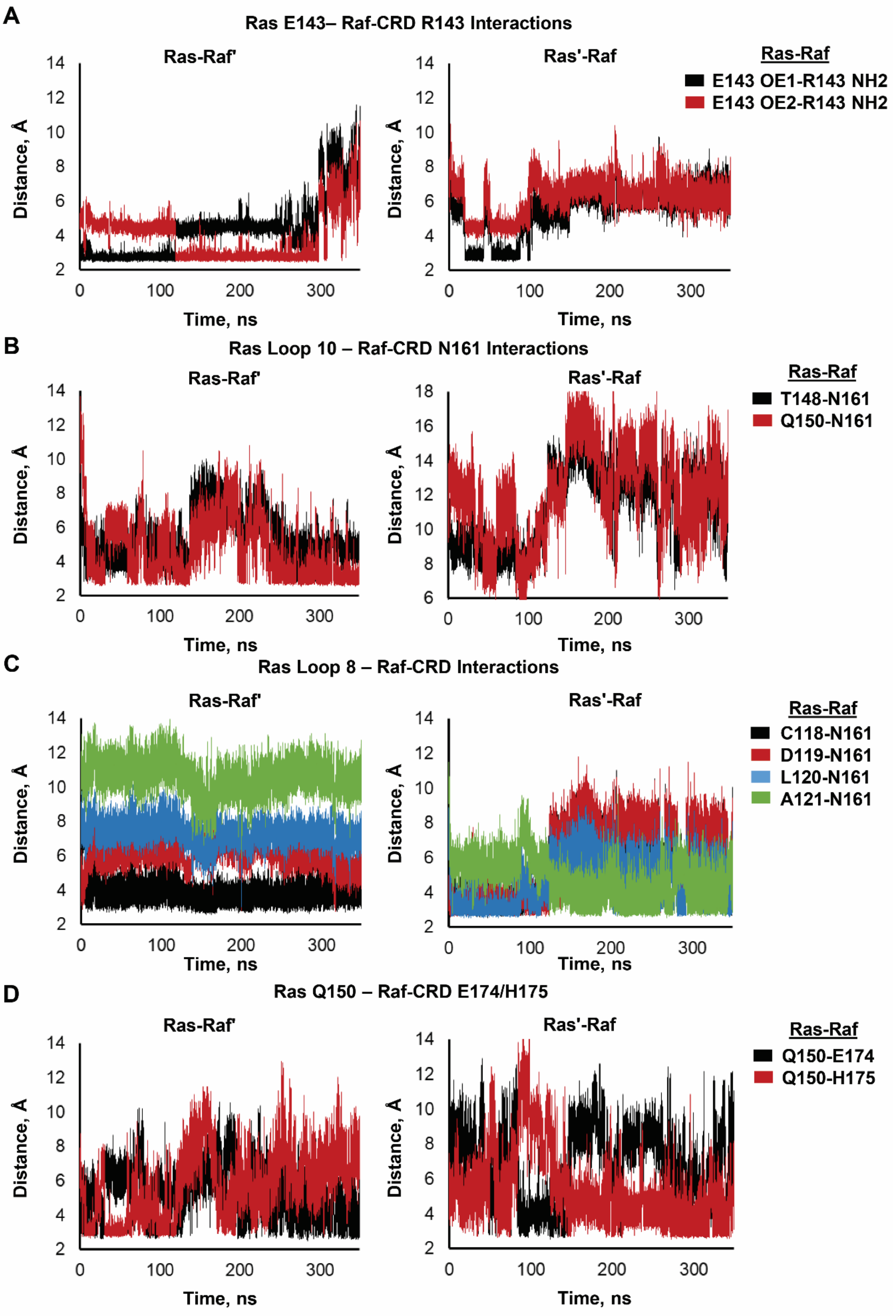
A
Ras E143– Raf-CRD R143 Interactions
Ras-Raf'
Ras'-Raf
Ras-Raf
E143 OE1-R143 NH2
E143 OE2-R143 NH2
Distance, A
Time, ns
B
Ras Loop 10 – Raf-CRD N161 Interactions
Ras-Raf'
Ras'-Raf
Ras-Raf
T148-N161
Q150-N161
Distance, A
Time, ns
C
Ras Loop 8 – Raf-CRD Interactions
Ras-Raf'
Ras'-Raf
Ras-Raf
C118-N161
D119-N161
L120-N161
A121-N161
Distance, A
Time, ns
D
Ras Q150 – Raf-CRD E174/H175
Ras-Raf'
Ras'-Raf
Ras-Raf
Q150-E174
Q150-H175
Distance, A
Time, ns

**Fig. S4. Ras/Raf-CRD' and Ras'/Raf-CRD interactions over 250 ns simulation time of the HRas/CRaf-RBD_CRD dimer complex (replicate 2).** . **(A)** Distances calculated between the sidechain of Raf-CRD residue R143 and the side chain of residue E143 of the opposing Ras protomer. Ras/Raf-CRD' interactions are shown on the left and Ras'/Raf-CRD interactions are shown on the right. **(B)** Distances calculated between the sidechain of Raf-CRD residue N161 and the side chains of T148 (black) and Q150 (red) located in loop 10 of the opposing Ras protomer. **(C)** Distances calculated between the sidechain of Raf-CRD residue N161 and the carbonyl backbone of residues C118 (black), D119 (red), L120 (blue), and A121 (green) in the opposing Ras molecule. **(D)** Distances calculated between the sidechains of Raf-CRD residues E174 (black) and H175 (red) and the side chain of residue Q150 of the opposing Ras protomer.

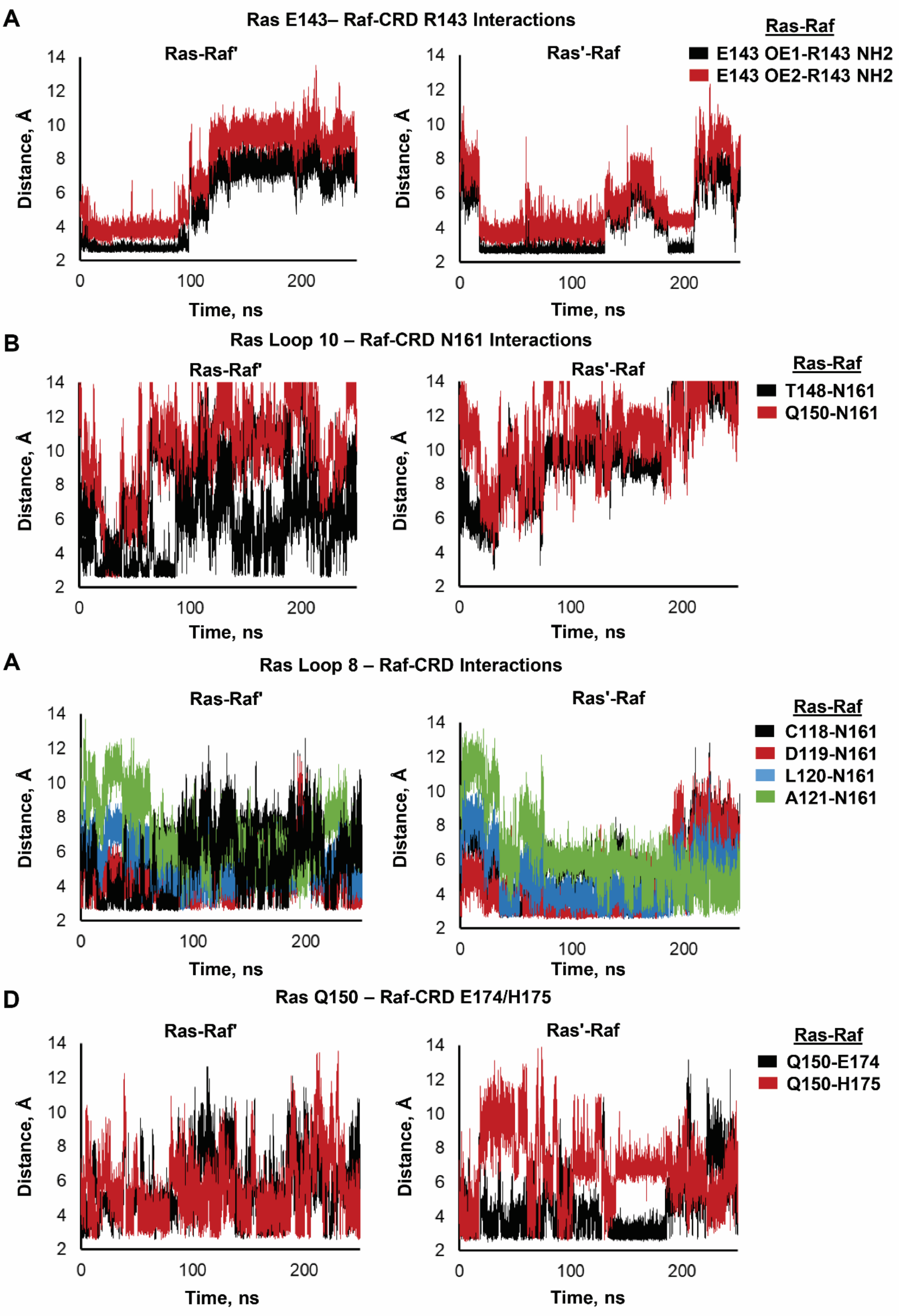
A
Ras E143– Raf-CRD R143 Interactions
Ras-Raf'
Ras'-Raf
Ras-Raf
E143 OE1-R143 NH2
E143 OE2-R143 NH2
B
Ras Loop 10 – Raf-CRD N161 Interactions
Ras-Raf'
Ras'-Raf
Ras-Raf
T148-N161
Q150-N161
A
Ras Loop 8 – Raf-CRD Interactions
Ras-Raf'
Ras'-Raf
Ras-Raf
C118-N161
D119-N161
L120-N161
A121-N161
D
Ras Q150 – Raf-CRD E174/H175
Ras-Raf'
Ras'-Raf
Ras-Raf
Q150-E174
Q150-H175
Distance, Å
Time, ns
0
100
200
2
4
6
8
10
12
14

**Fig. S5. Ras/Raf-CRD' and Ras'/Raf-CRD interactions over 250 ns simulation time of the HRas/CRaf-RBD_CRD dimer complex (replicate 3). (A)** Distances calculated between the sidechain of Raf-CRD residue R143 and the side chain of residue E143 of the opposing Ras protomer. Ras/Raf-CRD' interactions are shown on the left and Ras'/Raf-CRD interactions are shown on the right. **(B)** Distances calculated between the sidechain of Raf-CRD residue N161 and the side chains of T148 (black) and Q150 (red) located in loop 10 of the opposing Ras protomer. **(C)** Distances calculated between the sidechain of Raf-CRD residue N161 and the carbonyl backbone of residues C118 (black), D119 (red), L120 (blue), and A121 (green) in the opposing Ras molecule. **(D)** Distances calculated between the sidechains of Raf-CRD residues E174 (black) and H175 (red) and the side chain of residue Q150 of the opposing Ras protomer. Sequence alignment of BRaf and CRaf conserved region 1 containing the Raf-RBD and CRD.

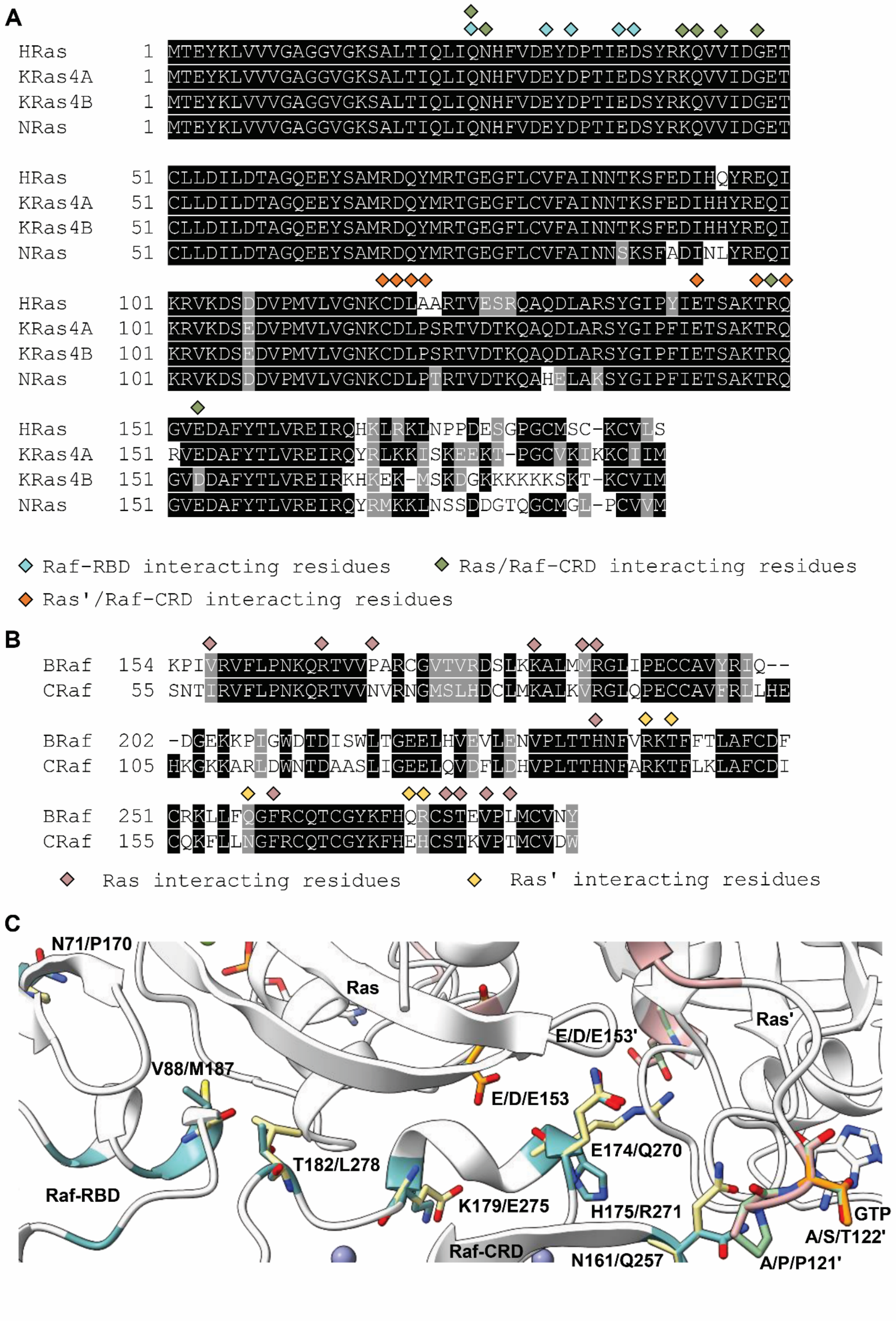
A
HRas 1 MTEYKLVVVGAGGVGKSALTIQLIQNHFVDEYDPTIEDSYRKQVVIDGET
KRas4A 1 MTEYKLVVVGAGGVGKSALTIQLIQNHFVDEYDPTIEDSYRKQVVIDGET
KRas4B 1 MTEYKLVVVGAGGVGKSALTIQLIQNHFVDEYDPTIEDSYRKQVVIDGET
NRas 1 MTEYKLVVVGAGGVGKSALTIQLIQNHFVDEYDPTIEDSYRKQVVIDGET
HRas 51 CLLDILDTAGQEEYSAMRDQYMRTGEGFLCVFAINNTKSFEDIHQYREQI
KRas4A 51 CLLDILDTAGQEEYSAMRDQYMRTGEGFLCVFAINNTKSFEDIHHYREQI
KRas4B 51 CLLDILDTAGQEEYSAMRDQYMRTGEGFLCVFAINNTKSFEDIHHYREQI
NRas 51 CLLDILDTAGQEEYSAMRDQYMRTGEGFLCVFAINNSKSFADINLYREQI
HRas 101 KRVKDSDDVPMVLVGNKCDLAARTVESRQAQDLARSYGIPYIETSAKTRQ
KRas4A 101 KRVKDSEDVPMVLVGNKCDLPSRTVDTKQAQDLARSYGIPFIETSAKTRQ
KRas4B 101 KRVKDSEDVPMVLVGNKCDLPSRTVDTKQAQDLARSYGIPFIETSAKTRQ
NRas 101 KRVKDSDDVPMVLVGNKCDLPTRTVDTKQAHELAKSYGIPFIETSAKTRQ
HRas 151 GVEDAFYTLVREIRQHKLRKLNPPDESGPGCMSC-KCVLS
KRas4A 151 RVEDAFYTLVREIRQYRLKKISKEEKT-PGCVKIKKCIIM
KRas4B 151 GVDDAFYTLVREIRKHKEK-MSKDGKKKKKKSKT-KCVIM
NRas 151 GVEDAFYTLVREIRQYRMKKLNSSDDGTQGCMGL-PCVVM
Raf-RBD interacting residues
Ras/Raf-CRD interacting residues
Ras'/Raf-CRD interacting residues
B
BRaf 154 KPIVRVFLPNKQRTVVPARCGVTVRDSLKKALMMRGLIPECCAVYRIQ--
CRaf 55 SNTIRVFLPNKQRTVVNVRNGMSLHDCLMKALKVRGLQPECCAVFRLLHE
BRaf 202 -DGEKKPIGWDTDISWLTGEELHVEVLENVPLTTHNFVRKTFFTLAFCDF
CRaf 105 HKGKKARLDWNTDAASLIGEELQVDFLDHVPLTTHNFARKTFLKLAFCDI
BRaf 251 CRKLLFQGFRCQTCGYKFHQRCSTEVPLMCVNY
CRaf 155 CQKFLLNGFRCQTCGYKFHEHCSTKVPTMCVDW
Ras interacting residues
Ras' interacting residues
C
N71/P170
Ras
Ras'
E/D/E153'
V88/M187
E/D/E153
E174/Q270
T182/L278
Raf-RBD
K179/E275
H175/R271
GTP
A/S/T122'
Raf-CRD
N161/Q257
A/P/P121'

**Fig. S6. Ras and Raf isoform specific residues at the Ras/Raf interface. (A)** Sequence alignment of the four Ras isoforms (HRas, KRas4A, KRas4B, and NRas) with residues denoted with a diamond corresponding to those that interact with the Raf-RBD (blue), with the Raf-CRD through the loop 3 interface (green), and those that interact with the Raf-CRD through the loop 8 interface (orange). **(B)** Sequence alignment of BRaf and CRaf conserved region 1 containing the Raf-RBD and CRD. Residues denoted with a pink diamond are involved in interactions at the Ras loop 3 / Raf-CRD interface and residues denoted with a yellow diamond are involved in interactions at the Ras' loop 8 / Raf-CRD interface. **(C)** Close up view of the Ras/Raf-CRD interfaces with isoform specific residue sidechains shown for CRaf (blue), BRaf (yellow), HRas (pink), KRas (green), and NRas (orange).

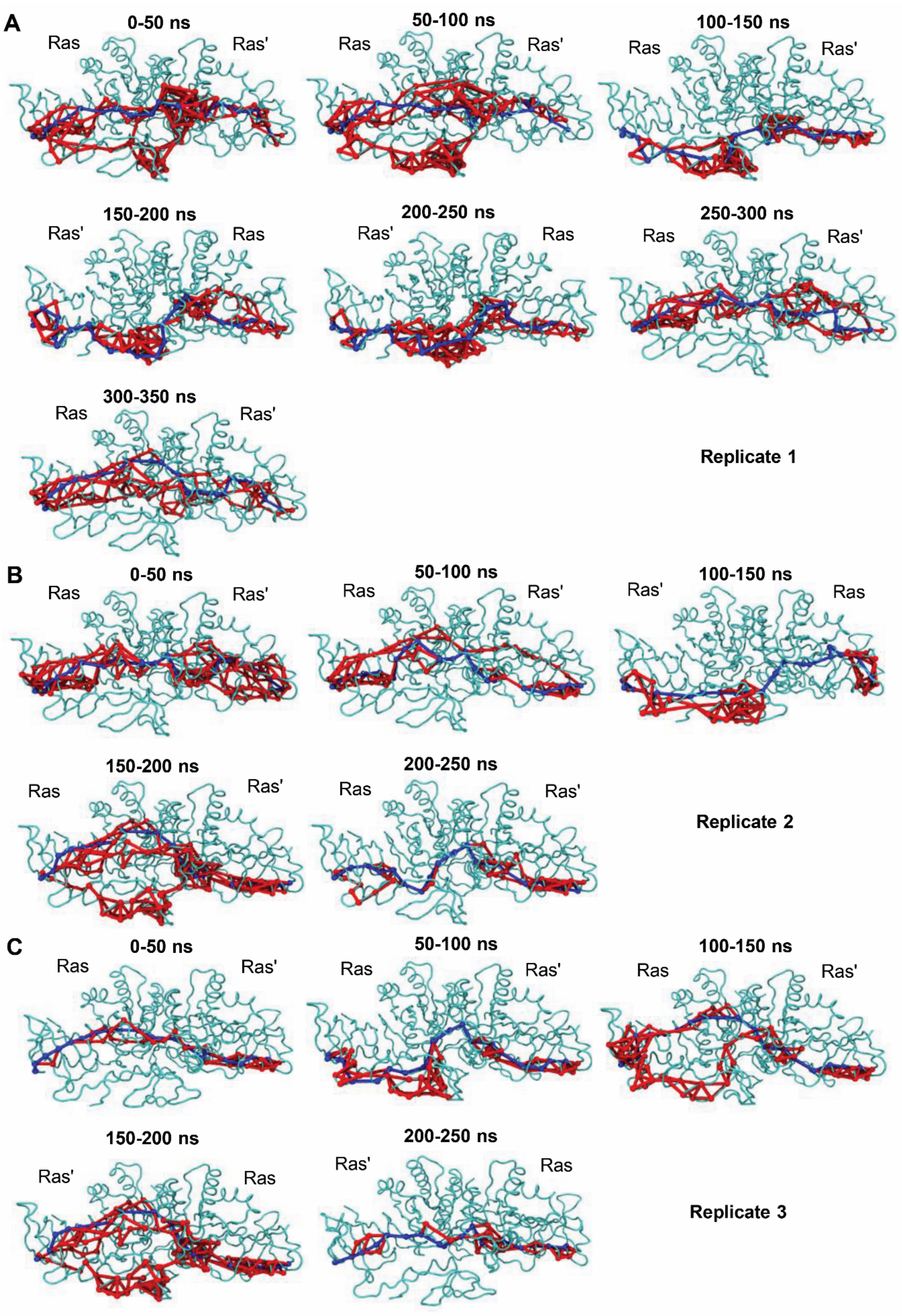
A
0-50 ns
Ras
Ras'
50-100 ns
Ras
Ras'
100-150 ns
Ras
Ras'
150-200 ns
Ras'
Ras
200-250 ns
Ras'
Ras
250-300 ns
Ras
Ras'
300-350 ns
Ras
Ras'
Replicate 1
B
0-50 ns
Ras
Ras'
50-100 ns
Ras
Ras'
100-150 ns
Ras'
Ras
150-200 ns
Ras
Ras'
200-250 ns
Ras
Ras'
Replicate 2
C
0-50 ns
Ras
Ras'
50-100 ns
Ras
Ras'
100-150 ns
Ras
Ras'
150-200 ns
Ras'
Ras
200-250 ns
Ras'
Ras
Replicate 3

**Fig. S7. Network view of all optimal and suboptimal path calculations between Raf-RBD residues D113 and D113'. (A-C)** Visualization of paths calculated for 50 ns segments of 350, 250, 250 ns total simulation time for the three replicates of the HRas/CRaf-RBD_CRD dimer modelled utilizing crystallographic symmetry showing the 4 modes of intermolecular information transfer involving helix 5, loop 3, and loop 8 regions of the Ras G-domain and the Raf-CRD. The optimal path, which corresponds to that with the least number of nodes, is shown in blue. All other paths are designated as suboptimal paths and are shown in red.
